# Cooperative mechanosensitivity and allostery of focal adhesion clusters

**DOI:** 10.1101/175273

**Authors:** D. C. W. Foo, E. M. Terentjev

## Abstract

We analyse an Ising-like Hamiltonian describing the free energy of cell adhesion on a substrate as a lattice of 3-state mechanosensing sites involving focal adhesion kinase (FAK). We use Monte Carlo stochastic algorithm to find equilibrium configurations of these mechanosensors in two representative geometries: on a 1D ring representing the rim of a cell on flat surface, and a 2D bounded surface representing the whole area of cell contact with flat surface. The level of FAK activation depend on the pulling force applied to the individual FAK-integrin via actin-myosin contractile networks, and the details of the coupling between individual sensors in a cluster. Strong coupling is shown to make the FAK sensors experience a sharp on-off behaviour in their activation, while at low coupling the activation/autoinhibition transition occurs over a broad range of pulling force. We find that the activation/autoinhibition transition of FAK in the 2D system with strong coupling occurs with a hysteresis, the width of which depends on the rate of change of force. The effect of introducing a mediating protein (such as Src) in limited quantity to control FAK activation is explored, and visualizations of clustering in both topologies are presented. In particular the results on the bounded 2D surface indicate that clustering of active FAK occurs preferentially at the boundary, in agreement with experimental observations of focal adhesions in cells.

## 1. Introduction

Cell adhesion plays an important role in many different physiological situations, including wound healing, cell migration, and development and the maintenance of tissues [1]. The structurally defined adhesion sites between cells and the extracellular matrix (ECM), that would eventually be called focal adhesions (FA), were first identified over 40 years ago [2], and characterized using electron and interference-reflection microscopy [2, 3, 4, 5]. Since then, immense progress has been made in elucidating their biochemistry and over 50 proteins have been identified as playing a role in the FA function; some of the details of their interactions remain open questions [6, 7]. Of particular importance is a 120 kDa tyrosine-kinase protein now called focal adhesion kinase (FAK), whose phosphorylation and spatial distribution were linked with the presence of transmembrane integrins and appropriate extracellular ligands [8, 9, 10] and which has since been identified as a central node in the FA signalling network [11, 12].

FAK has three main domains: the N-terminal 4.1 protein, erzin, radixin, moesin (FERM) domain, the kinase domain in the middle, and the C-terminal focal adhesion targeting (FAT) domain, through which the FAK is localized to FA via integrin associated proteins paxillin and talin [13, 14], see Fig. 1. The FERM domain has been implicated both as an autoinhibitor of the kinase domain and as having some role in the kinase domain activation [15]. More recent studies have solidified the FERM domain autoinhibitory role, but have attributed the proper activation of the kinase domain to a separate protein, Src [16, 17]. In its native ground state, the FERM and kinase domains are held in close proximity by physical bonds, blocking access to phosphorylation sites on the kinase domain. In this paper, this will be called the closed configuration, [c], see Fig. 2. In order to activate the kinase, the FERM domain needs to be pried apart from the kinase domain to allow autophosphorylation of FAK’s Tyr397 residue. This will be called the open state, [o]. It is possible for FAK to spontaneously close again after phosphorylation of Tyr397 [13]. Src uses the phosphorylated Tyr397 as an anchorage point to bind to FAK to block the autoinhibition and complete the activation of the kinase domain through phosphorylation of other tyrosine residues in the kinase domain, resulting in the active state [a]. Once fully activated, FAK is unable to close without first dephosphorylating the kinase domains [16] and so transitions between these three states must be done in sequence: [c] ⇌ [o] ⇌ [a].

Much of the literature has focused on the biochemistry of FA, but more recently considerable evidence has built up pointing to the roles of mechanical force [18, 19, 20] and substrate elasticity [21, 22, 23, 24, 25] in regulation of cell behaviour, with work done on both single sensors [26, 27] and clusters of molecules [28, 29, 30, 31]. Here we follow in the vein of recent work on conformational spread and allostery in proteins, motivated by clustering of membrane receptors in bacterial chemotaxis [32, 33, 34]. It was shown that interactions (coupling) between sensors lead to the formation of sensor clusters, onset of binary (switch-like) behaviour and enhanced sensitivity. We analyse a simplified scheme of FA cluster formation using FAK as the central player, and show how cluster formation may be regulated by the application of mechanical force.

**Figure 1.**
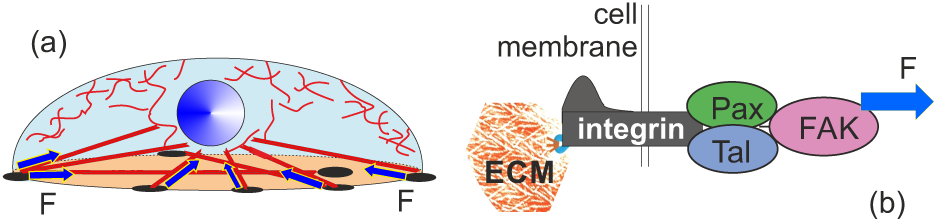
Cell adhesion on a substrate. Integrins spanning the cell membrane attach to biochemical markers on the ECM outside the cell, and to paxillin and talin inside the cell, which themselves bind to FAK. The cytoskeleton exerts a pulling force on this complex. Focal adhesions form predominantly (although not uniquely) on the rim of the flat surface.

We consider the FAK, possibly as part of some larger complex, connected to integrin at the C-terminal via paxillin or talin and to the actin cytoskeleton at the N-terminal via the actin nucleator Arp2/3 [35]. The actin cytoskeleton exerts a pulling force *F* generated by the actin-myosin contractile network [36]. Initially, these integrin-FAK-actin sequences are homogenously strewn around the cell. We then consider two cell topologies; first, a 1D ring of 1000 sites with cyclic boundary conditions, which is meant to emulate the edge of a cell on a flat substrate, as in Fig. 1, where FA are noted to predominantly form. Secondly, we consider a 2D square lattice truncated to form a bounded, approximately circular surface of 1009 sites, which is meant to model the entire adhesion surface underneath the cell on a flat substrate, on which it will be shown that interactions between mechanosensing complexes naturally lead to clustering on the boundary of the surface. For the sake of clarity, we repeat that while reference will repeatedly be made to FAK alone, this is a shorthand for the machanosensing complex as a whole, which may involve many other proteins and we are considering a greatly simplified model thereof.

## 2. Methods

### 2.1. Single mechanosensor: three-state model

As mentioned before, FAK are strongly associated with transmembrane integrins via connecting proteins such as paxillin or talin [13]. They may thus be considered bound to their ECM sites despite being always contained within the interior of the cell. Due to their high population, the simplest model to describe collective FAK behaviour and clustering is thus a discrete model, where the membrane is partitioned into a collection of sites on which a single FAK molecule is placed (or equivalently a single mechanosensing complex of integrin, paxillin, talin, FAK and whatever other proteins may be pertinent). Each site *i* then holds two Ising (binary) variables, *s*_*i*_*, r*_*i*_ *∈* {*-*1, 1} which collectively encode the conformation of the FAK at that site as shown in Table 1.

**Figure 2:**
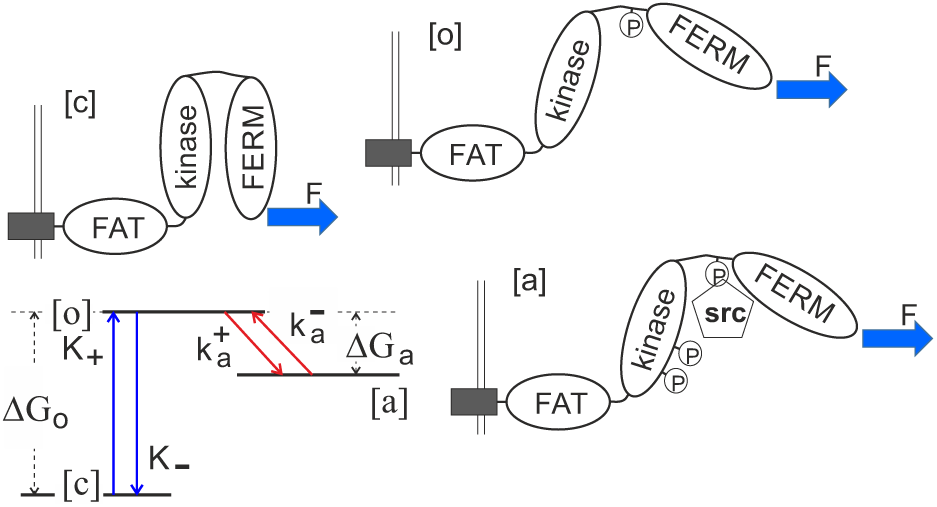
Three conformations of FAK. The FAT domain indirectly binds to integrins that bridge the cell membrane and in turn connect to the ECM. The cytoskeletal pulling force *F* is applied to the FERM domain via actin-myosin contractile network. The closed ground state [c] has the lowest free energy, the open metastable state [o] allows phosphorylation of Tyr397. The open conformation of FAK has its kinase ‘activated’ by binding Src, [a], which also prevents it from directly returning back to the native state. The corresponding map of the threestate system has its energy levels and transition rates marked.

**Table 1.** Values of the binary variables *s*_*i*_*, r*_*i*_ in the three conformational states of FAK.

| Conformation at $i$ | $s_i$ | $r_i$ |
| --- | --- | --- |
| Closed | +1 | -1 |
| Open | -1 | -1 |
| Active | -1 | +1 |

The conformation of a FAK molecule is then described by the Hamiltonian

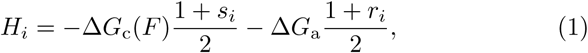

where *H*_*i*_ is the energy of a single non-interacting mechanosensor at site *i* and Δ*G*_c,a_ are the energy differences between the conformations (see Fig. 2), defined positive when the open conformation has the highest energy. It is implicit that the free energy level of the open conformation is zero, so that the free energy of the closed conformation varies linearly with the force experienced at a particular site, via the mechanical work contribution:

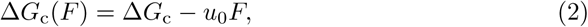

where *F* is the average force experienced by the cell as a whole, and *u*_0_*≈*0.3 nm is a characteristic length of the physical bond in FERM-kinase (which controls the [c] ⇌ [o] transitions), on the order of the size of a single amino acid residue.

The energy density for a collection of *N* noninteracting FAK molecular sensors is then straightforwardly given by

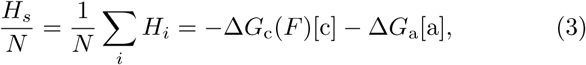

where *H*_*s*_ is the total energy of all non-interacting sensors. We have here introduced the ‘order parameters’

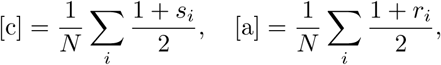

which represent the fractions of closed and active FAK, respectively. The proportion of open FAK is not independent: [o] = 1–[a]–[c], wherein we make the assumption that the total amount of FAK remains constant in time. In the absence of interactions between neighbouring FAK, the Hamiltonian *H*_*s*_ leads to a force-dependent level of equilibrium activation, [a], as shown in Fig. 3 below. Regarding the specific value of Δ*G*_c_, The MD simulation study [37] estimated the value as Δ*G*_c_ *≈* 28.5*k*_*B*_*T*, but it was noted that this value appears to be too high, given the low-affinity nature of interdomain hydrophobic interactions in proteins. Measurements made on a different multidomain protein gave a value of *≈*11*k*_*B*_*T* [38], but this was for only 2 hydrophobic contacts while FAK is known to have more [37]. In this paper, we have gone with an intermediate value of Δ*G*_c_ = 15*k*_*B*_*T*, in keeping with the recent study of FAK morphology[27], both to have some grounding in reality and to ensure convergence of simulations in a timely fashion. The related estimate of the energy change associated with activation was chosen to be Δ*G*_a_ = 5*k*_*B*_*T*, a typical value for phosphorylation of a protein. Staying in units of energy, the force was scaled by the factor *k*_*B*_*T /u*_0_, which for *T* = 36*°*C, is approximately equal to 14 pN.

### 2.2. Coupling of mechanosensors: Ising-like model

Interactions between individual sensors (integrin-FAK-actin sequences) are introduced through an interaction Hamiltonian *H*_*J*_, the details of which depend on how the 3 FAK conformations affect the energies of neighbouring molecules. In a simple Ising-like model where the variables {*s*} and {*r*} take binary values, nearest neighbours interact to raise or lower the system energy by an amount *J* depending on the value of the site variables. In our 3-state system, there are potentially 6 different interactions between nearest neighbours. However, in order to limit the space of possibilities, given the conformational dissimilarity between closed FAK and the two open forms, we have limited our discussion to schemes where [c]-[o] and [c][a] type neighbours do not interact at all. Examples of interaction schemes and the reduction of energy in multiples of the parameter *J* are illustrated in Table 2

**Table 2:** Several distinct coupling schemes are listed, to guide the analysis below. The values in the table show, in units of *J*, how much the overall free energy is reduced when different FAK conformations are in the neighbouring sites.

| Coupling scheme | c-c | c-o | c-a | o-o | o-a | a-a |
| --- | --- | --- | --- | --- | --- | --- |
| a/o equal | 0.5 | 0 | 0 | 1 | 1 | 1 |
| o-a weak | 0.5 | 0 | 0 | 1 | 0.5 | 1 |
| o-o strong | 0.5 | 0 | 0 | 1.5 | 1 | 1 |
| c-c = a/o | 1 | 0 | 0 | 1 | 1 | 1 |

For example, the interaction scheme ‘c-c = a/o’ has the nearest neighbour interaction Hamiltonian taking the familiar Ising form:

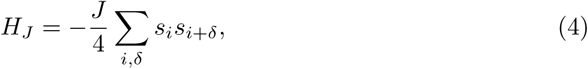

where an unimportant constant term has been omitted. The interaction between similar (dissimilar) *s*_*i*_ decrease (increase) the energy by the amount of *J/*2, with a further division by 2 to account for double-counting in pairs of neighbours. The interaction Hamiltonian for the ‘o-a weak’ scheme takes a slightly more complicated form:

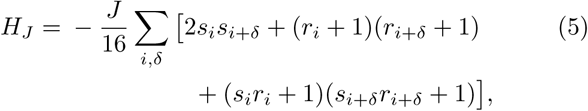

which is not separable in *s* and *r*. It is thus not immediately evident that conclusions drawn from analysis of the standard Ising model hold in this case. In both examples, the index *i* runs over all sites and the index *δ* runs over the nearest neighbours of each *i* (which are two in the 1D ring sensor configuration, and four in the 2D area configuration, except near the edge).

### 2.3. Monte Carlo simulation

Numerical results were obtained via Monte Carlo simulations utilising the Glauber algorithm to determine transition probabilities between states. Two different cell topologies were simulated: the 1D ring with 1000 mechanosensing sites, and the 2D finite square lattice with boundary conditions applied to form quasi-circular surface with 1009 sites. Each site can adopt 3 states, closed, open and active, which can only be accessed sequentially as described above. The cells were initialised with all sites closed and with zero force applied.

At every step of the simulation, a site was randomly selected and a partition function calculated for the cell configurations accessible by at most one state transition of the designated site, keeping in mind which state transitions are allowed and which are forbidden. For example, if the randomly selected site was active, the partition function was calculated using the energies of the present cell macrostate, and the macrostate that is identical except that the selected site is in the open state. The partition function calculated was then used to determine the probabilities of transition and a random number generated to determine which transition occurs.

The simulation was then made to run for a billion steps in the 1D case and ten million in the 2D case, with macrostate data recorded every ten thousand or hundred steps for the last million or ten thousand steps, giving a hundred samples in total from which the average equilibrium activity of the cell was calculated. This constitutes a single reading of equilibrium activity at a particular value of applied force. Readings are then taken for force values from *u*_0_*F* = 0 to *u*_0_*F* = 20*k*_*B*_*T*.

The different values for the number of steps were chosen to balance accuracy and simulation runtime. In the 1D case, no phase transition and thus no hysteresis was expected in analogy with the classic Ising model, so the number of steps was lengthened till the results reflected this. In the 2D case, a phase transition was expected which in Monte Carlo simulations should manifest as hysteresis, so a smaller value was used to speed up the simulation as the 2D simulation inherently ran slower than the 1D one. The effect of increasing or decreasing the number of steps in the 2D case is discussed in greater detail in the Results section.

It should be noted here that unlike the more commonly used Metroplis algorithm, the Glauber algorithm allows for ‘null’ transitions where the system macrostate does not change even though the present macrostate is a local maximum in energy. For example, if the randomly selected site is in the open state, there is a finite probability that it will remain so after the Monte Carlo step even if zero force is applied to the cell. The Glauber algorithm was chosen over the Metropolis algorithm because it admits a more natural generalisation to systems with more than two states and because it has a physical origin of a system interacting with a heat bath, as opposed to the the more mathematical origin of the Metropolis criterion [39].

### 2.4. Slope and hysteresis width

The maximum slope of the various activation-force graphs of the 1D ring was found by approximating the crossover region as a straight line and finding the slope between the first point beyond 20% and the last point before 80% of the maximum activation. This approximate method was chosen to smooth out the effect of fluctuations that become more pronounced when using methods that rely on taking the differentials between successive points.

The hysteresis width in the case of the 2D surface was found by using the lever rule to interpolate the force at half maximum activation and finding the full width at this half maximum value. The rate of change of force was altered by changing the number of simulation timesteps per fixed change of applied force.

### 2.5. Monte Carlo with limited Src

Later simulations, in which the quantity of Src was limited, were performed by modifying the probability of [o]*→*[a] transition in proportion to the amount of remaining Src. That is, the partition function was calculated as normal but the probability of an open to active transition was multiplied by the fraction of free Src. In this way the initial simulated results represent the condition of Src in excess. While reference is made to Src specifically, the results obtained should be regarded in generality, considering a limited quantity of any accessory protein necessary for stabilising activation of a mechanosensing site, giving the cell another method of controlling the overall level of activation.

## 3. Results and discussion

### 3.1. Sensors distributed on 1D ring (rim of the cell)

Figure 3 indicates that at zero pulling force, most of the FAK molecules are in the closed (native folded) state. At high force, almost no FAK molecules remain in the closed state, [o]+[a]*→*1; in this regime, the equilibrium ratio between [o] and [a] is determined by the free energy of Src-mediated phosphorylation: [o]/[a]= exp[−Δ*G*_a_*/k*_*B*_*T*], which sets the maximum possible level for [a], which for Δ*G*_a_ = 5*k*_*B*_*T* gives [a]_max_ = 0.9933. Note that, since we take the scaled free energy difference to be Δ*G*_c_*/k*_*B*_*T* = 15, it is around the matching value of scaled force *F u*_0_*/k*_*B*_*T* = 15 that the closed (folded) state becomes statistically rare. It is interesting to note that the numerical data for the equilibrium fraction of non-closed states [o]+[a], in Fig. 3 can be fitted very closely by the ‘standard’ sigmoidal function (1 + tanh[(*F - F*_*∗*_)*/w*])*/*2, with the mid-point at *F*_*∗*_ *≈* 10*k*_*B*_*T /u*_0_ and the width of the transition *w ≈* 1*k*_*B*_*T /u*_0_.

The effect of different coupling schemes of mechanosensor interaction is readily described using mean field theory. Although the Ising model on a 1D ring has an exact analytical solution [40], the meanfield approach is simpler and gives perhaps a better insight into the limiting behaviour, despite erroneously predicting a phase transition in 1D. The *s*_*i*_*s*_*i*+*δ*_ term causes an effective lowering of the energy of the closed conformation and thus a shift in Δ*G*_c_ of

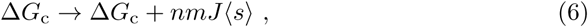

with *n* the number of nearest neighbours, *m* the multiplier value found in the scheme Table 2, and *(s)* is the mean value of the Ising (binary) variable *s*_*i*_. Similarly terms involving (*s*_*i*_ + 1)(*s*_*i*+*δ*_ + 1) cause shifts of the form

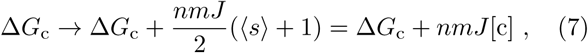

and the analogous terms involving *r*_*i*_ and *s*_*i*_*r*_*i*_ result in the lowering of the active, or the open energy levels, respectively. These effects are most visible in the limit of large force, where the energy of the closed conformation has been raised by the application of force (so [c] is no longer the ground energy state), and the average activity depends only on the energy difference between open and active conformations. Additional o-o coupling lowers the energy of the open conformation, reducing Δ*G*_a_ and inhibiting activation. Strengthened c-c coupling causes a rightward shift of the transition curve, as in Fig. 4, making the transition occur at higher pulling force, but does not affect the behaviour at high or low force, as expected. Tuning of the coupling scheme thus allows a cell to directly control the effective range of its protein sensors and their efficacy.

**Figure 3:**
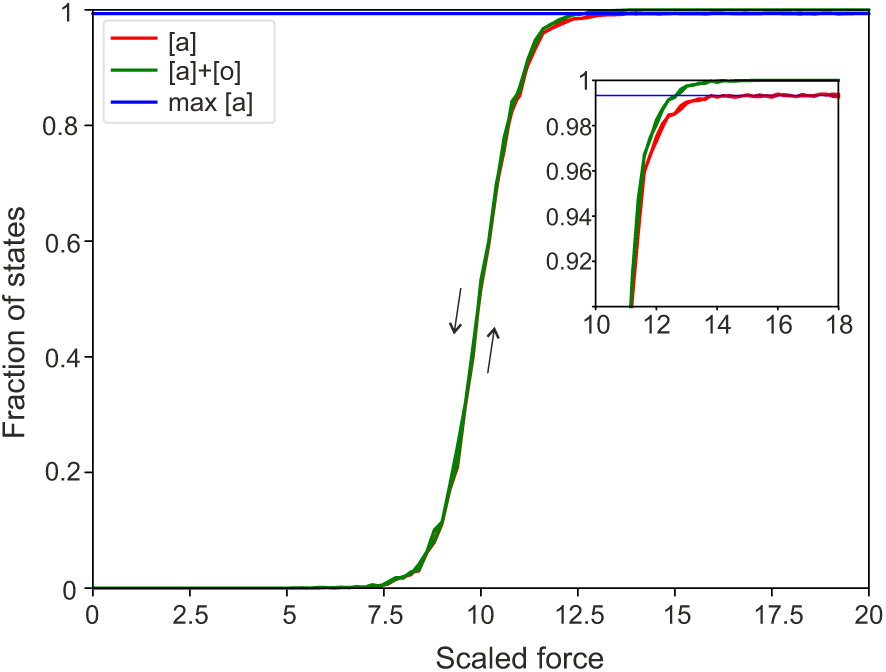
Force-dependence of activation in isolated FAK sensor. As force increases, the energy of the closed conformation is raised and the total fraction of FAK that is not closed ([o]+[a]) tends to unity. As activation is a purely biochemical process, the phosphoprylation barrier Δ*G*a does not depend on force, and in the high force limit the proportion of active FAK [a] tends to the value predicted by Boltzmann statistics: with Δ*G*_a_ = 5*k*_*B*_*T*, [a]_max_ = 0.9933. Both increasing*F* and decreasing-*F* curves are shown, appearing on top of each other, thus indicating that equilibrium has been achieved. The inset shows a more expanded region near saturation.

**Figure 4:**
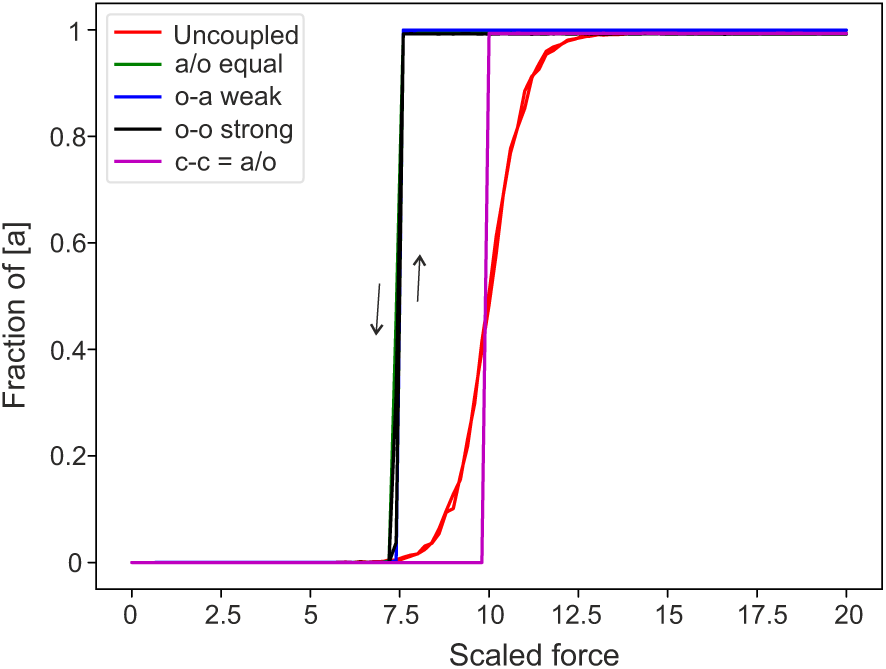
Activation of a ring for different coupling schemes. The position of transition from completely closed to mostly active, on increasing pulling force, is shown not to depend strongly on the coupling scheme used, with the general nature of the transition remaining the same. In particular, the conclusion of no first order phase transition at finite temperature holds the same as for the 1D Ising model. The coupling parameter *J* = 5*k*_*B*_*T*. The ‘uncoupled’ curve is the same as in Fig. 3.

Physically, the interaction Hamiltonian may arise directly from protein-protein interactions between hydrophobic or hydrophilic groups. Alternatively it may be indirectly mediated by deformations of the membrane caused by the presence of FAK [41] or other membrane-associated proteins. As values for the interaction strengths are not currently known, the ‘o-a weak’ scheme was chosen to be the subject of more extensive analysis, on the basis that the closed conformation has fewer active sites and less surface area exposed, decreasing the relative strength of c-c coupling compared to o-o and a-a, and the presence of Src, likewise, results in a slight mismatch between open and active conformations.

The different coupling schemes were tested using the method described in the previous section on the 1D ring configuration, which is meant to represent the rim of the cell attached to a flat substrate, with integrin-FAK sensors distributed along this rim. The result of varying coupling scheme is shown in Fig. 4 while the result of varying coupling strength in a fixed scheme is shown in Fig. 5. The position and shape of the transition are shown not to vary considerably under the conditions simulated, which can be explained using the mean field theory described above. In the limit of high force, only the effective free energy difference between open and active states matters (since all [c] states are gone), which is why the ‘uncoupled’, ‘a/o equal’, and ‘c-c = a/o’ schemes tend to the same plateau value of [a]*→*1*/*(1+exp[*-*Δ*G*_a_*/k*_*B*_*T*]) as these schemes do not have any differences between o-o, oa or a-a interactions. As previously mentioned, the strengthened c-c coupling in the ‘c-c = a/o’ scheme, as opposed to the ‘a/o equal’ scheme, causes an effective drop in the energy of the closed state, resulting in a rightward shift of the transition, since a greater force is required to overcome this effective reduction of energy.

**Figure 5:**
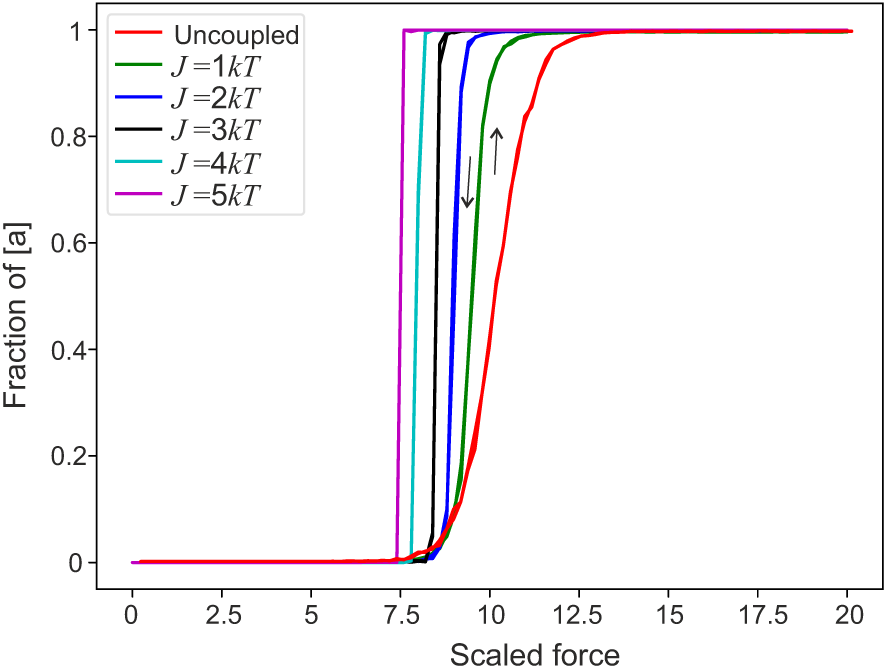
Activation of a ring for different coupling strengths. As the strength of coupling *J* increases, the activation transition becomes both sharper (i.e. occurring over a more narrow range of force), and more pronounced, achieving a higher proportion of activation at any given force. As in previous plots, both increasing-*F* and decreasing-*F* data are plotted, with curves on top of each other indicating equilibrium conditions.

**Figure 6:**
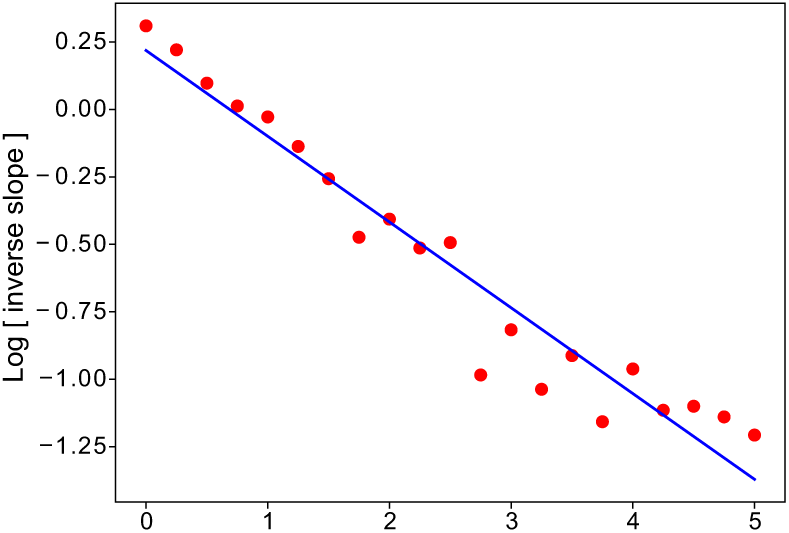
Inverse slope of the crossover region for FAK activation. The slope of switchover increases exponentially with the coupling strength *J*. Red dots show raw data collected from many simulations of a ring with varying values of *J*. The line of best fit is: 1.2*·*exp[*-*0.32*J/k*_*B*_*T*], as emphasized by the log-linear plot.

Additional o-o coupling in the ‘o-o strong’ scheme results in a lowering of the open energy level, resulting in a reduced overall level of activation of [a] *→* 1*/*(1 + exp[*-*(Δ*G*_a_*-J* [o])*/k*_*B*_*T*]) where we have inserted the values *n* = 2 for a 1D ring and *m* = 1*/*2 for oo coupling *J* /2 stronger than o-a or a-a coupling. However, given the exponential dependence on the coupling strength, this reduction is not readily visible. Combining this expression with the constraint on FAK proportions in the high force limit, [o] + [a] = 1 gives a transcendental equation for either [a] or [o]. For the ‘o-a weak’ scheme, terms involving both *r*_*i*_ and *s*_*i*_*r*_*i*_ in the interaction Hamiltonian *H*_*J*_ result in both the open and active levels shifting. However, at high force, very few FAK units are still closed and so *s*_*i*_ is almost always equal to *-*1; therefore, while the *r*_*i*_ term lowers the energy of the active level, the *s*_*i*_*r*_*i*_ term, or more accurately the *s*_*i*_*r*_*i*_ + 1 term, becomes very small leading to only a small reduction in the energy of the open state, in turn, leading to a very high level of activation of [a] *→* 1*/*(1 + exp[*-*(Δ*G*_a_ + *J* ([a] *-* [o]))*/k*_*B*_*T*]). In all coupling schemes, however, there is a general trend that, compared to the ‘uncoupled’ sensors ensemble, the force-mediated transition from the predominantly closed (native, folded FAK) to predominantly open/active FAK is sharper. As expected in the 1D topology, the transition is still continuous, with the curves for increasing and decreasing force overlapping within simulation error. This, combined with the genera visual similarity of the curves gives us some freedom to select a favoured scheme for analysis, in the absence of relevant data from real cells, with the hope that general conclusions will be unaffected by the choice of scheme. As previously mentioned, we will look at the ‘o-a weak’ scheme as such a characteristic example.

Increasing the coupling parameter *J* in the ‘o-a weak’ scheme was found to lead to increased activation in the high force limit, as expected from mean field theory, since the gap between the energy levels of open and active states widens. Unfortunately, this is hard to see in Fig. 5; what is clear however is that the transition occurs at lower force for higher *J*, which could not be predicted from mean field theory as the theory breaks down in the transition region for 1D. Nevertheless, this can be understood in terms of how the averages *⟨s⟩*, *⟨r⟩* and *⟨sr⟩* evolve under force: at low force, *⟨s⟩ ≈* 1, *⟨r⟩ ≈-* 1 and *⟨sr⟩ ≈* −1, so the energy of the closed state is lowered, while the energies of the open and active states are unchanged. As force increases, ⟨*s⟩* decreases and ⟨*r⟩* increases, but because these changes must happen in sequence (on a particular site, *s*_*i*_ *→*− 1 before *r*_*i*_ *→* 1), *⟨sr⟩* remains at approximately −1. Physically, the open state is still the highest energy state and so remains mostly unpopulated. Thus at intermediate force, the open energy remains unchanged while the closed energy level rises both due to force and the mean-field effect of coupling from the decrease in ⟨*s⟩*, while the energy of the active state falls due to a similar mean-field effect. The magnitude of the mean field effects depend linearly on *J*, so at higher *J* the transition to open occurs at a lower force.

**Figure 7.**
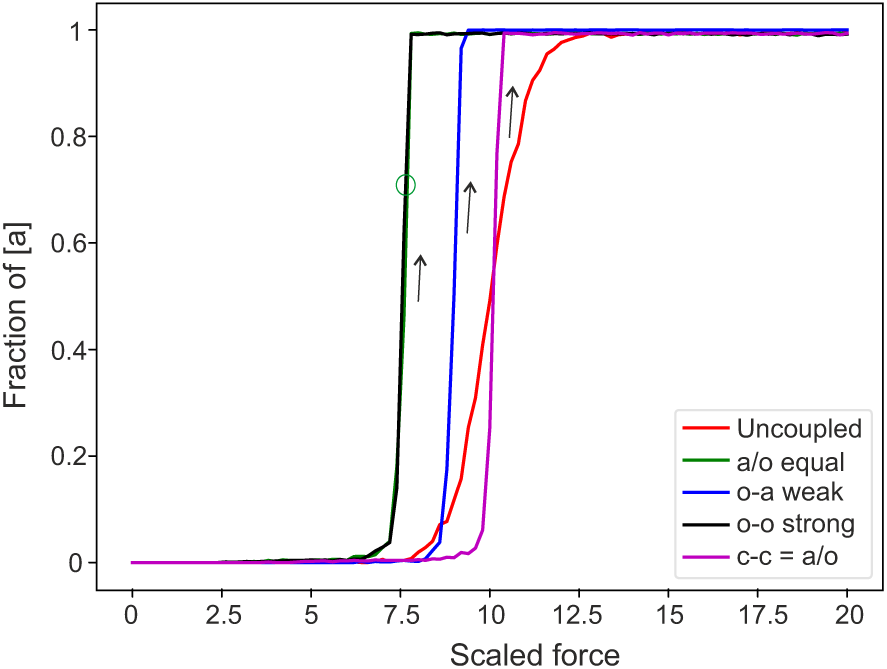
Activation of a surface for different coupling schemes. For clarity, only the increasing-*F* behaviour is shown here (the hysteresis is discussed in Fig. 8: in 2D case the curves for decreasing force did not coincide; the widths of hysteresis depended on the coupling scheme). The limiting behaviour at high force is similar to that seen in the 1D ring. The coupling parameter *J* = 2.5*k*_*B*_*T*.

The width of the transition region was found to decrease exponentially with increasing *J*, see Fig. 6. In this illustration, we characterise the transition width by the slope of the [a] vs. force plots in Fig. 5, so that the high slope represents the narrow (sharp) transition.

### 3.2. Sensors spread on 2D bounded surface

The same analysis was conducted on the 2D bounded surface. All qualitative results obtained from the 1D system were expected to carry over to 2D, as indeed seen in Fig. 7 where the high force activation levels are the same as in 1D (to achieve the same values, the coupling constant *J* has been halved to make up for the doubled number of nearest neighbours). Edge effects were not visible. However, unlike in 1D, the 2D system is expected to exhibit a discontinuity in the evolution of activation against force. This prediction comes from analysis of the Ising model in 2 and higher dimensions. At zero field, as temperature is decreased, the Ising model exhibits a first order phase transition from paramagnetic to ferromagnetic at a critical temperature. Below the critical temperature *T*_*c*_, the Ising order parameter, *m*, takes a value of *±m*_0_ as the symmetry is spontaneously broken. Turning the field on while below the critical temperature thus selects either +*m*_0_ or *-m*_0_ as the true ground state depending on the sign of the field, with a discontinuous switch at zero field. In our model, force is analogous to field and coupling strength *J* is varied instead of *T*. Since the contribution of coupling to the effective field is proportional to *J/k*_*B*_*T*, the condition *T < T*_*c*_ becomes *J > J*_*c*_, thus at high *J* a discontinuity in [a] is expected as force is varied.

**Figure 8.**
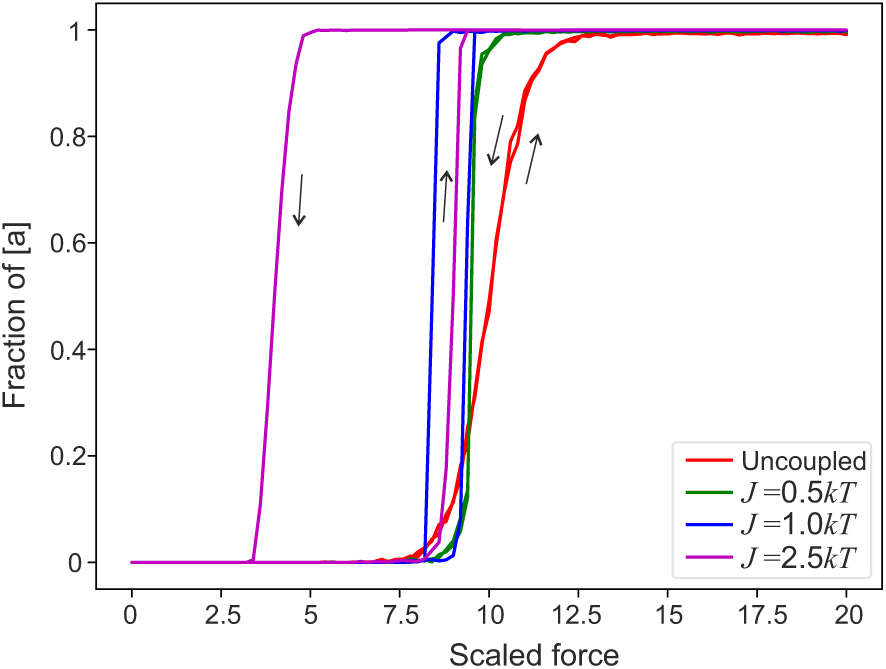
Activation of a surface for different coupling strengths. At zero or low coupling strength, the equilibrium distribution was achieved. However, above a critical value of coupling parameter *J*_*c*_ *<* 0.5*k*_*B*_*T* the system develops a discontinuity in the‘order parameter’ [a]. As the simulation only applies a small thermal perturbation, this manifests as hysteretic behaviour, since the system only seeks out a local minimum and does not have sufficient time to find the true thermodynamic equilibrium at any given rate of force ramp.

No discontinuity was observed in simulations as the magnitude of thermal fluctuation emulated by the chosen algorithm was insufficient to allow the system to fully explore the parameter space in the allotted number of steps (time of equilibration at each value of pulling force). Instead, hysteresis curves were obtained for coupling above a certain critical value. This is shown in Fig. 8 where hysteresis is not evident for *J* = 0 or *J* = 0.5*k*_*B*_*T*, but is rapidly increasing for *J* = *k*_*B*_*T* and above. Onsager’s exact analysis of the Ising model <u>i</u>n 2D [42] gives the critical coupling 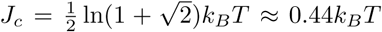, although it bears mentioning that his and our definition of *J* are not the same: in the standard formulation of the Ising model, *J* is half the energy difference between like and unlike pairs of sites in the absence of applied field, while in our model *J* is merely a base parameter modified by the multipliers in the table of coupling schemes (Table 2) to find the reduction in system energy when adjacent sites have some combination of states. As our chosen coupling scheme involves two coupled Ising variables, the direct translation of the classical Onsager result into our language is nontrivial. Qualitatively, however, it is clear that some critical value for the coupling must exist. In a real cell, the presence of hysteresis leading to ‘delaying’ the activation / autoinhibition transition on changing of force, may represent a beneficial feature of collective mechanosensing processes as it provides a measure of stability against environmental fluctuations.

As the hysteresis loop is not an equilibrium phenomenon, its width must depend on the rate at which the force is changed. This is explored in Fig. 9, where the width of the hysteresis decreases with a decreasing rate of change of force, appearing to tend to some value Δ*F*_hyst_ *<* 3*k*_*B*_*T /u*_0_. This limiting quasi-equilibrium value is a result of the small thermal fluctuation driving state evolution of the system, not being enough to escape from a local minimum of the potential energy landscape in the alloted time. If the fluctuations were large enough to allow the system to find the global minimum, or if the number of timesteps were increased sufficiently, there would be a thermodynamically equilibrium first-order transition, with a sharp discontinuity of [a] at a certain *J* dependent critical force, with no hysteresis. As the rate of change of force is non-vanishing, the system retains more memory of its previous configuration; mostly closed for increasing force and mostly active for decreasing force, widening the hysteresis loop as the system does not have time to seek out the metastable state described by the local energy minimum and instead remains on the trailing edge of a moving potential well. This widening of hysteresis due to the system response lagging behind the applied force is evident even in 1D if the rate of change of force is high [43].

**Figure 9:**
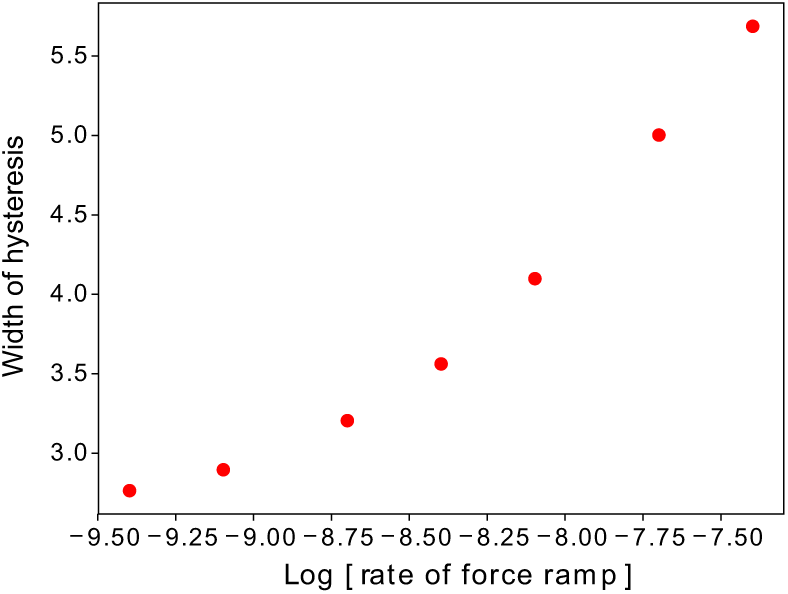
Width of the hysteresis loop. We take the transition curves with the highest hysteresis in Fig. 8, at the coupling parameter *J* = 2.5*k*_*B*_*T*; the rate of force is measured by the (scaled) ratio Δ*F/*Δ*t* between incremental steps. At slow rate of change of force the system has more time to find and settle into the closest local minimum, but not enough to find the global minimum. At higher rates of force ramping, the system response starts to lag behind the applied force, resulting in widening of the hysteresis loop.

### 3.3. Limited supply of Src

So far analysis has assumed that the concentration of Src in the cell is not appreciably changed in the course of mechanosensing, and so the transition between open and active conformations is completely governed by the energy change between the open and active configurations. This leads to the cell being unable to regulate the proportion of active FAK, as this is controlled only by the magnitude of intracellular force, which is itself a response to the stiffness of the substrate. The cell cannot easily alter the coupling strengths between FAK mechanosensing sites either as this is a function of the biochemistry, which can only be altered on evolutionary timescales and not the lifetime of a single cell. To regulate FAK activity and clustering then, the cell has to either control the amount of FAK itself, or the amount of accessory protein necessary for transduction of the signal of the active FAK sites. We focus here on the protein Src, a 60 kDa tyrosine kinase that activates the kinase domain of FAK [16, 7, 9, 13, 14]. Being a smaller protein, its concentration can be altered much faster than the concentration of FAK but we are here using it symbolically for any molecule that can similarly affect the signal transduction between FAK and the cytoskeletal actin network, and thus regulate intracellular forces, clustering of FAK, and the formation of FAs.

We limit the amount of Src in the system by considering a fixed number of total Src that may be either bound to FAK or free in the cytosol, altering the probability of transition from the open to active state by the proportion of free Src remaining. This not only puts a hard ceiling on the maximum proportion of active FAK, but also makes highly active configurations less likely. Results on the 1D ring and 2D surface topologies are shown in Figs. 10 and 11 respectively. The amount of Src is expressed as a proportion normalised to the amount of FAK, that is the number of sites in the system, so for example [Src] = 0.4 in the ring topology of 1000 FAK sensors means 400 Src molecules available in total. In both 1D and 2D, the effect of limiting Src is shown to suppress the high-force plateau level of activation of the cell to a level below [Src], as expected. In 1D, the width of transition is also affected, getting wider as Src becomes more limited. This is because each activation has a greater impact on the probabilities of subsequent activations if [Src] is low, resulting in more gradual curves. However, the onset of activation is roughly constant at *F ≈* 7.5*k*_*B*_*T /u*_0_ as this point is only determined by when the force is sufficient to trigger transitions from the closed state to the open state and is unaffected by the presence or absence of Src. Once some FAK open, the initial probability of activation is the same and it is only later in the evolution of the system that the effect of limited Src becomes important.

In 2D, the right edges of the hysteresis loops, where force is increasing, coincide. While the point of onset of activation is expected to coincide as it does in 1D, the fact that the slope is also unaffected indicates that the system is not evolving adiabatically and the rate of force increase is too high, causing the system to be carried along the trailing wall of a moving potential energy barrier. As a result the decreasing slope of switchover seen in 1D is not observed. In contrast, the left edges, where force is decreasing, do not coincide, with the width of the hysteresis loop decreasing with decreasing [Src]. In the high-force limit, the system essentially takes the form of an Ising ferromagnet in an applied field, with a balance struck between minimising the number of domain walls between open and active clusters while operating under a constraint on the number of sites that may align with the field. With low [Src], many domain walls already exist and it is relatively easy to allow the system to revert to a mostly open, and then mostly closed, state by allowing transitions along the already extant domain walls. As [Src] increases, the number of domain walls in the fully active state decreases, introducing an additional energy barrier as the system must not only transit to the higher energy open state to settle into the closed ground state but also overcome the cost of creating domain walls. This causes the fully active state to persist longer and require a greater reduction of force to trigger the formation of domain walls and eventual complete deactivation of the system.

**Figure 10:**
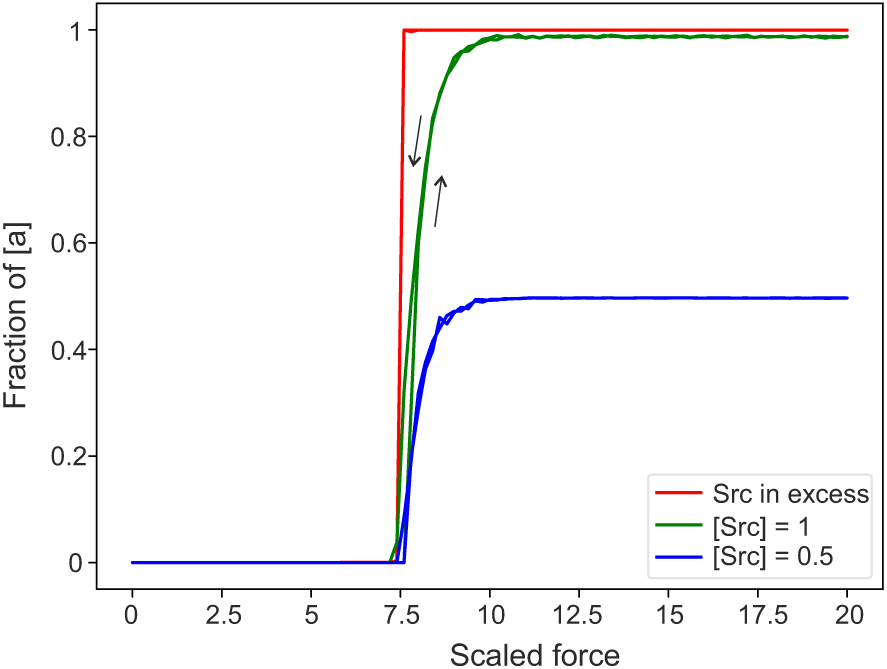
Equilibrium activation of a 1D ring for different concentrations of Src. Decreasing [Src] reduces the high-force level of activation and makes the crossover regime less sharp. The coupling parameter *J* = 5*k*_*B*_*T*.

**Figure 11:**
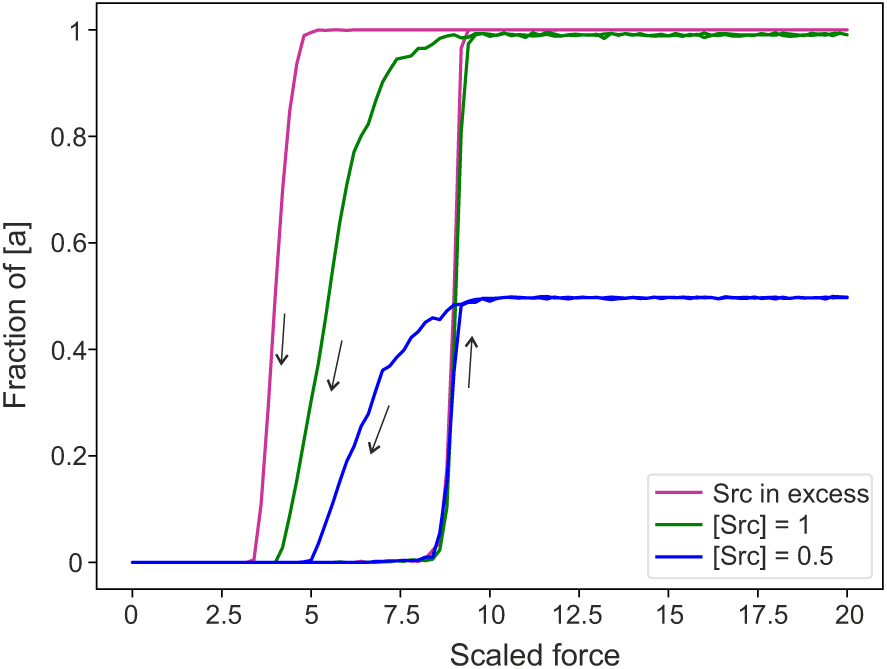
Activation of a 2D surface for different concentrations of Src. Decreasing [Src] reduces the high-force level of activation, and also the width of the hysteresis loops, while the position of the activation transition remains is fixed. The coupling parameter *J* = 2.5*k*_*B*_*T*.

### 3.4. Clustering of sensors on rings and surfaces

The results obtained so far have tracked the value of the average order parameter [a], which while pertinent to the biological activity of the cell, does not in and of itself reflect clustering. Yet, the FAK sensor units are expected to self-organise in clusters to some extent as doing so lowers the system energy for the same level of activation. Figs. 12 and 13 confirm that expectation, showing that in both topologies, at a sufficiently high force, a non-zero coupling *J* works to organise the active FAK units into clusters, compared to the uncoupled case *J* = 0 which has a random assortment of open and active FAK at proportions set by the interplay of FAK energies and limited Src. In all cases, the limit of [Src] = 0.5 was chosen to allow a sufficient proportion of [o] states, and make clustering evident. The presented snapshots of the system were taken after one million steps of the chosen algorithm, giving each site approximately one thousand transition opportunities. At *F* = 20*k*_*B*_*T /u*_0_, the [c]*→*[o] and the first [o]*→*[a] transition probability is *e*^5^*/*(1 + *e*^5^) *≈* 0.9933 so a thousand steps should be sufficient to achieve a representative snapshot of equilibrium behaviour. To confirm this, the system was allowed to continue running a hundred times longer to a hundred million steps and no detectable change in morphology was evident.

While the 1D system merely shows a collection of active clusters of varying sizes assembled on the rim of model cell, the 2D system gives a physical significance to the edge: the boundary of the cell surface in contact with the ECM. We see that large active clusters are always connected to the boundary; in the given snapshot for *J* = 2*k*_*B*_*T*, the largest contiguous cluster not adjacent to the cell boundary, whether open or active, contains only a single site. This is because the sites on the boundary, having fewer nearest neighbours and thus fewer potentially stabilising pair interactions, are more susceptible to changes of state, setting up the boundary as a nucleating surface for clusters of different states. At the top right of the snapshot we see a fairly large cluster of open sites having nucleated from within an active cluster, itself also connected to the boundary. While the number of sites is small compared to the number of molecules within a typical cell, the results nevertheless emphasise that interactions are necessary for self-assembly into clusters, and provide an explanation for why FAs in real cells form almost exclusively on the edge of their contact surfaces.

**Figure 12:**
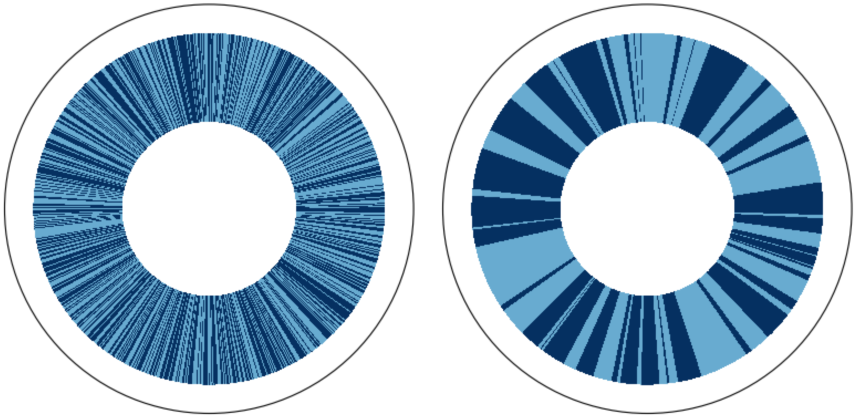
Role of sensor coupling in forming clusters of activated sensors. Snapshot of a 1D ring under high force (scaled) *F* = 20*k*_*B*_*T /u*_0_, and [Src]= 0.5, comparing the cases without and with coupling. The ‘equilibrium’ after 1 million steps. Light blue segments are open FAK, and dark blue are active FAK positions; white portions are closed FAK positions (almost none remain at this high force). Starting from an initial state of all FAK closed, coupling promotes FAK clustering and suppresses the proportion of closed states by increasing the closed state energy.

**Figure 13:**
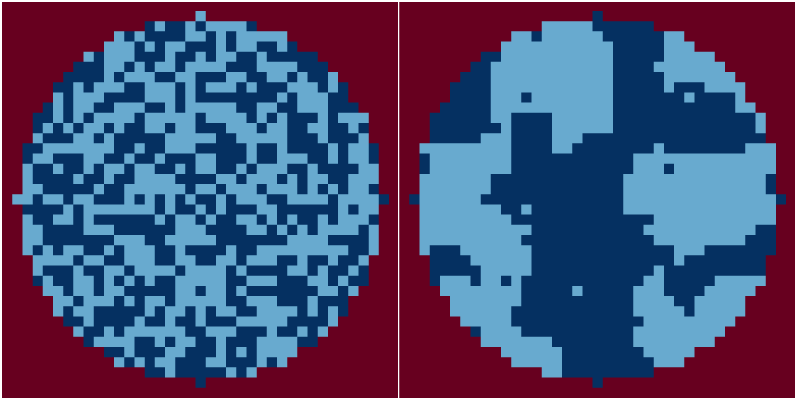
Role of sensor coupling in forming clusters of activated sensors. Snapshot of a bounded surface under high force (scaled) *F* = 20*k*_*B*_*T /u*_0_, [Src]= 0.5, after 1 million steps. Light blue squares are open FAK, and dark blue are active FAK positions within the cell footprint; white portions are closed FAK positions (almost none remain at this high force). Starting from an initial state of all FAK closed, coupling promotes clustering of active FAK and causes it to occur preferentially close to the boundary, due to the decreased number of interactions setting up a nucleating surface. Coupling also causes the closed state to be disfavoured through increasing of the closed state energy.

## 4. Conclusions

The recent work on FAK mechanosensors [27] concentrated on isolated protein complexes, their activity regulated via complex feedback pathways that ultimately result in a rapid approach to an equilibrium level of intracellular force dependent only on substrate stiffness, assuming parameters describing cell biochemistry are known [26, 44]. In other words, the intracellular force is insensitive to the initial conditions of the cell and settles on the value prescribed by the environment. Here we did not examine the effect of substrate stiffness, merely explored the fact that substrate stiffness directly maps onto the pulling cytoskeletal force [27]. In this way we simply utilised the tilting of potential energy’ as a result of increasing force to initiate the transition in FAK activation, which leads to a change of round cells on soft substrates and widely-spread star-shaped cells on stiff substrates, when focal adhesions and stress fibers dominate the cell morphology.

Our analysis implies that models of mechanosensors in which only the average activity of cooperative proteins is examined could miss many of the important aspects of the dynamics of transitions between conformational states. Cooperative sensors might be even better at filtering fluctuations and coping with high pulling forces that the cell generates on stiffer substrates. The coupling between the neighbouring sensor conformations results in a positive feedback.

Here we examined the collective configuration of FAK mechanosensors (obviously, in a complex with integrins and other mediating proteins), taking the intracellular cytoskeletal force as a variable and considering the effect of allosteric interactions between FAK sites. Monte Carlo simulations were run in order to find equilibrium (steady state) conformations on 1D ring (representing mechanosensors exclusively on the edge of cell adhesion surface), and 2D bounded surface topology (representing the sensors distributed homogeneously everywhere on the adhesion footprint). The results show that the fraction of activated FAK depends on the coupling scheme, the coupling strength and the amount of mediating protein, Src. The presence of a mediating protein in particular was noted to give the cell a degree of control over the maximum value of activated FAK, and thus over cluster formation. A similar effect is seen in, for example, bacterial chemotaxis receptors [34] or the insulin-mediated clustering of epidermal growth factor [45, 46]. Non-equilibrium (rate-dependent) behaviour was explored in the 2D system, with hysteresis evident at all tested rates of change of force, owing to the inevitable thermal fluctuations present in our the simulation, as well as in the real system. Clustering of sensors (representing the formation of focal adhesions) was observed in both topologies, with the 2D system in particular exhibiting preferential nucleation of clusters on the boundary, in agreement with the observed preferred distribution of focal along the edges of adherent cells on stiff substrates.

The hysteresis of mechanosensor response is particularly interesting. This effect only appears in a 2D surface topology, and only when the strength of allosteric coupling between sensors is high enough; if the sensors acted independently – or if they were restricted only to an open rim of the cell contact surface, the sensor response is fully reversible: on any given substrate, the level of cytoskeletal force has a definitive relation with the level of active FAK. But the hysteresis that occurs for strongly coupled sensors interacting in the whole plane of adhesion adds much stability against against force fluctuations: once the threshold force is reached and FAK sensors activate, this level of signal (which is regulated by Src) remains stable even if the force decreases below this threshold. Only a significant drop in pulling force would allow FAK units to auto-inhibit in this interacting environment. Perhaps this is the reason why in practice the focal adhesions are found everywhere on the surface on contact, even though more frequently near the open rim of the cell where lamellipodia develop.

## Acknowledgments

The authors have benefited from extensive discussions with K. Chalut, D. Bray and S. Bell. This work has been funded by UROP scheme by the Department of Physics, University of Cambridge.

